# Dynamic product-precursor relationships underlie cuticular lipid accumulation on maize silks

**DOI:** 10.1101/2021.03.31.437946

**Authors:** Keting Chen, Liza E. Alexander, Umnia Mahgoub, Yozo Okazaki, Yasuhiro Higashi, Kouji Takano, Derek Loneman, Tesia S. Dennison, Miriam Lopez, Reid Claussen, Layton Peddicord, Kazuki Saito, Nick Lauter, Karin S. Dorman, Basil J. Nikolau, Marna D. Yandeau-Nelson

## Abstract

The hydrophobic cuticle is the first line of defense between aerial portions of a plant and the external environment. On maize silks, the cuticular cutin matrix is infused with cuticular lipids, consisting of a homologous series of very-long-chain fatty acids (VLCFAs), aldehydes, and hydrocarbons that serve as precursors, intermediates, and end-products of the elongation, reduction, and decarbonylation reactions of the hydrocarbon-producing pathway. To deconvolute the potentially confounding impacts of the silk microenvironment and silk development on the hydrocarbon-producing pathway, spatio-temporal cuticular lipid profiling was conducted on the agronomically important inbreds B73 and Mo17, and their reciprocal hybrids. Statistical interrogation via multivariate analyses of the metabolite abundances of the hydrocarbon-producing pathway demonstrate that the cellular VLCFA pool is positively correlated with the cuticular lipid metabolome, and this metabolome is primarily affected by the silk microenvironment and the plant genotype. Moreover, genotype has a major effect on the pathway, with increased cuticular hydrocarbon and concomitant reduction of cuticular VLCFA accumulation on B73 silks, suggesting that conversion of VLCFAs to hydrocarbons is more effective in B73 than Mo17. Statistical modeling of the ratios between cuticular hydrocarbons and cuticular VLCFAs reveals the complexity of the product-precursor ratio relationship, demonstrating a significant role of precursor chain length. Longer-chain VLCFAs are preferentially utilized as precursors for hydrocarbon biosynthesis. Collectively, these findings demonstrate maize silks as an effective and novel system for dissection of the complex dynamics of cuticular lipid accumulation in plants.

**One-sentence Summary:** The product-precursor ratios in the cuticular hydrocarbon-producing pathway are impacted by fatty acid precursor chain length, plant genotype and the spatio-temporal dynamic gradients of maize silks.

## INTRODUCTION

The cuticle is the external hydrophobic barrier covering the epidermis of most aerial terrestrial plant organs, an exception being the bark of trees (Riederer and Schreiber, 2001). The cuticle limits transpirational water loss (Yeats and Rose, 2013) and thus has a role in protecting the organism from such abiotic stresses as drought, salinity, and temperature. Moreover, additional protective roles have been suggested, including protection from ultraviolet radiation (Krauss et al., 1997; Shepherd and Wynne Griffiths, 2006), and from biotic stresses, such as fungal and bacterial pathogens and herbivory by insects (Eigenbrode and Espelie, 1995; Serrano et al., 2014). In general, these extracellular cuticular lipids can include alkyl derivatives such as hydrocarbons, aldehydes, primary and secondary alcohols, ketones, and wax esters, which are metabolically derived from very long chain fatty acids (VLCFAs). Additional components include triterpene derivatives, such as beta-sitosterol, stigmasterol, lupeol and alpha- and beta-amyrins (Jetter et al., 2006). The specific composition of the cuticular lipid metabolome is dependent on the organism, tissue or organ, and temporal stage of tissue and organ development (Shepherd and Wynne Griffiths, 2006; Samuels et al., 2008). For example, in maize, the cuticular lipids on juvenile and adult leaves are primarily composed of alcohols, aldehydes and esters, with only about 1% and 17% being hydrocarbons, respectively (Bianchi et al., 1984; Bianchi et al., 1985). In contrast, for maize silks, which are the stigmatic floral tissues that receive pollen and facilitate fertilization of the ovule, cuticular lipids are particularly rich in hydrocarbons, comprising 40-90% of these lipids, with only minor amounts of VLCFAs, aldehydes and alcohols (Yang et al., 1992; Perera et al., 2010; Loneman et al., 2017; Dennison et al., 2019).

The favored model for hydrocarbon production in plants is via the reduction of a saturated or unsaturated VLCFA-CoA to an aldehyde intermediate, which appears to subsequently undergo decarbonylation to produce either a saturated (alkane) or unsaturated (alkene) hydrocarbon (Cheesbrough and Kolattukudy, 1984; Bernard et al., 2012; Jetter et al., 2018). Because the majority of VLCFAs are comprised of an even number of carbon atoms (*2n*, where *n* represents the number of 2-carbon units used to assemble the VLCFA), this process generates products with an odd-numbered carbon chain length (i.e., *2n – 1*). However, if the process begins with odd-numbered VLCFAs (i.e., *2n + 1* carbons), then even-numbered hydrocarbon products are produced (i.e., *(2n + 1) – 1 = 2n*). Such even-numbered hydrocarbons occur in many cuticles, including that of maize silks (Yang et al., 1992; Loneman et al., 2017; Dennison et al., 2019). This sequence of hydrocarbon-forming reactions can occur in parallel at each VLCFA-CoA chain length (i.e., at every value of *n*), thereby generating a homologous series of cuticular hydrocarbon products with alkyl chain lengths ranging from 19-33 carbon atoms. Importantly, the predicted VLCFA precursors, the aldehyde intermediates and the hydrocarbon products of this decarbonylation pathway are all present in the extractable silk cuticle (Loneman et al., 2017), permitting the assessment of relationships between these substrates, intermediates, and products. Indeed, based on the assessment of VLCFAs, aldehydes, alkanes, and alkenes collected from maize silks, a pathway model involving a series of parallel reactions has been proposed for the biosynthesis of both alkanes and alkenes (Perera et al., 2010).

As with all biological processes, this decarbonylative hydrocarbon biosynthesis pathway is genetically programmed and is regulated by many factors that integrate environmental and developmental cues. Indeed, both genetic and environmental factors impact the cuticular lipid metabolome on maize silks. For example, multiple characterizations of the maize silk cuticular lipid metabolome revealed substantial differences (i.e., up to 10-fold) in lipid abundance among various panels of inbred genotypes, demonstrating the breadth of natural variation in cuticular composition (Miller et al., 2003; Perera et al., 2010; Loneman et al., 2017; Dennison et al., 2019). Moreover, within any individual maize inbred line, cuticular lipid accumulation varies along the length of the silks. Specifically, cuticular hydrocarbons accumulate at up to five-fold higher levels on the portions of silks that have emerged from the protective husk leaves, as compared to the portions that are still encased by these husk leaves (Miller et al., 2003; Perera et al., 2010; Loneman et al., 2017; Dennison et al., 2019), demonstrating an impact of silk microenvironment on hydrocarbon biosynthesis. Thus, it is precisely the dynamic character of these natural systems that permits the statistical interrogation of the hydrocarbon-producing network among maize inbreds.

Herein, the hydrocarbon-producing network is characterized via cuticular lipid metabolome profiling of silks along a spatio-temporal gradient from two agronomically important inbred lines, B73 and Mo17, as well as their reciprocal hybrids. Supervised and unsupervised multivariate analyses of the cuticular lipid profiles along the silk length reveal that both microenvironment and genotype of the silks, but not silk development, greatly impact cuticular lipid compositional dynamics, particularly the relationships between hydrocarbon-products and VLCFA-precursors. Significantly, the product-precursor ratio varies according to the precursor carbon chain length, deepening our understanding of the factors impacting the reduction-decarbonylation pathway of hydrocarbon synthesis.

## RESULTS

To model the metabolic network for hydrocarbon production and deconvolute the potential impacts of development, microenvironment, and genetics on the dynamics of the cuticular lipid metabolome, we profiled extracellular cuticular lipids along the lengths of silks collected at 3-days post silk-emergence. These silks were deeply sampled (∼20 biological replicates) from four maize genotypes (inbred lines B73 and Mo17, and their reciprocal hybrids, B73×Mo17 and Mo17×B73) in the summer growing seasons of 2014 and 2015. The collected silks were dissected into five contiguous segments, named sections A-E, from base to tip (Figure 1A). These sections reflect the spatio-temporal gradient of the silk, and include the microenvironment transition of the silks, as sections A-C were husk-encased, and sections D and E were emerged into the external environment (McNinch et al., 2020). All aforementioned factors are analyzed with the 2014 data set, and the addition of the 2015 data allowed for the analysis of environmental effects. All cuticular lipid accumulation data is available in Supplemental Table S1.

**Figure 1.**
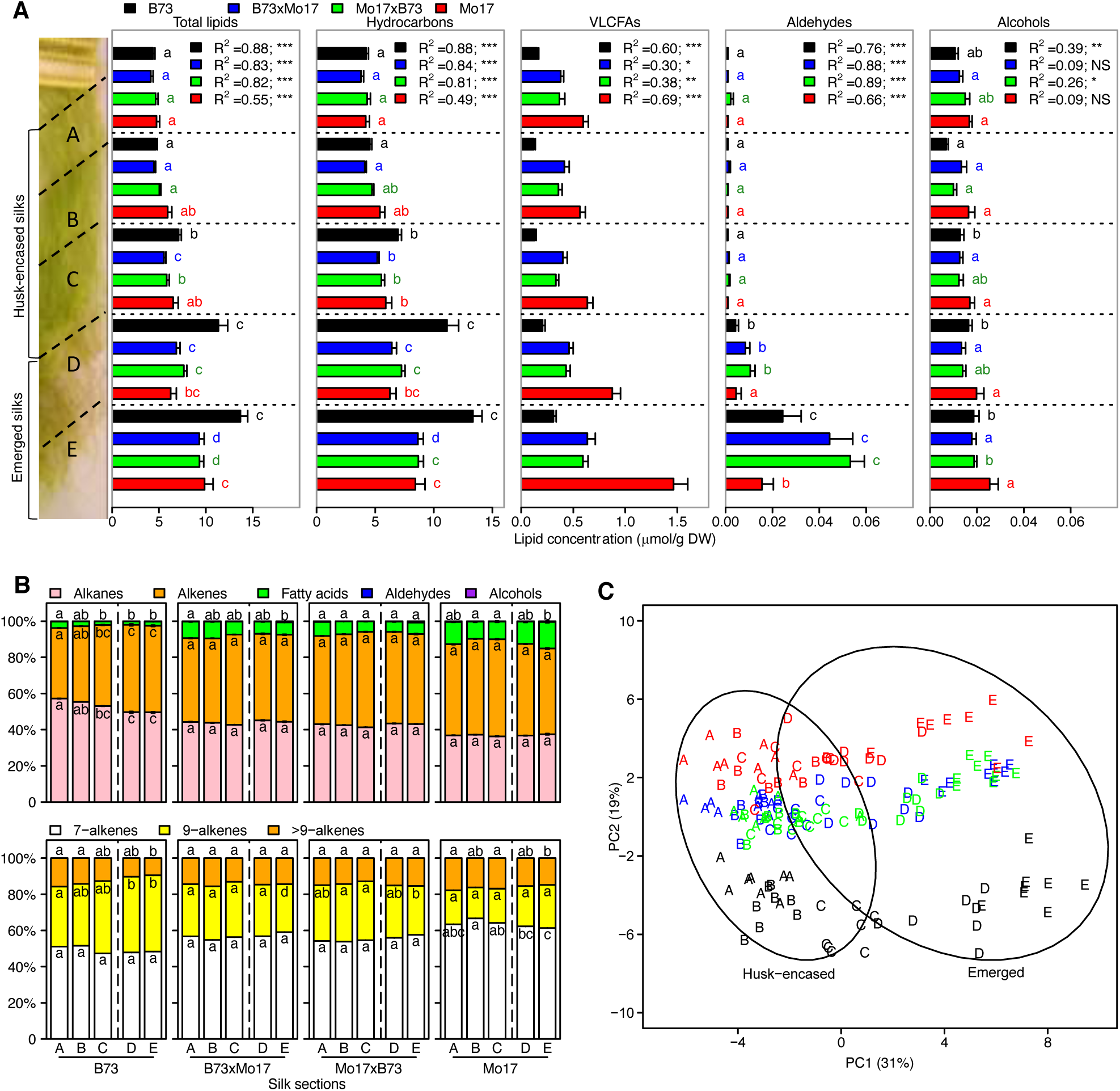
Spatio-temporal cuticular lipid profiles of maize silks from inbreds Mo17 and B73, and their reciprocal hybrids. **(A)** Concentrations of total cuticular lipids and individual lipid classes from silks harvested 3-days post-silk emergence and cut into five allometric sections, A-E (see Methods). The bold dashed line depicts the transition point between husk-encased (sections A, B and C) and emerged (sections D and E) portions of the silks. For each genotype, the changes in concentration of each lipid class along the A to E gradient were fitted to a quadratic regression model. The resulting R^2^ values are listed and the associated p-values are denoted by asterisks (***, p <0.0001; **, p <0.001; *, p <0.05). Different letters associated with data bars of the same color denote a statistically significant difference in accumulation between silk sections within a genotype (p <0.05; Tukey’s Honestly Significant Difference (HSD) test). Seven to eight replicates were evaluated per combination of genotype and silk section, constituting a total sample size of 158. **(B)** Relative compositions of cuticular lipids along the silk length. Proportion of each cuticular lipid class relative to total cuticular lipid accumulation. Aldehydes and alcohols comprise <1% of cuticular lipid metabolome and are not depicted in the figure. Alkenes are comprised predominantly of 7- and 9-monoene classes, and “>9” constituents that include 14- and 15-monoenes, and two dienes. Different letters within data bars for a given genotype denote a statistically significant difference in lipid relative abundance between silk sections for a given metabolite class (p<0.05; Tukey’s HSD test). **(C)** Principal component analysis (PCA) of cuticular lipid metabolite abundances among different silk sections. Each data point, which represents the concentrations of the 45 cuticular lipid metabolites profiled from each silk sample, is labeled by silk section (A-E) and color-coded according to genotype. Percentages represent the amounts of variance explained by the first and second principal components (PC1 and PC2). Ovals represent 95% confidence ellipses for emerged and husk-encased samples.

### Dynamic changes in the cuticular lipid metabolome along the silk length in different genotypes

Depending on genotype and silk section, hydrocarbons account for 85-97%, and VLCFAs account for 2-15% of total extractable cuticular lipids; whereas aldehydes and alcohols are detected only in trace amounts (<1%) (Figure 1, A and B; Supplemental Table S1). Irrespective of the genotype, total cuticular lipid load increases acropetally (section A to E; Figure 1). A three-fold increase in total cuticular lipid abundance is observed along this spatio-temporal gradient for inbred line B73, whereas total abundance only doubles for inbred line Mo17, and the two reciprocal hybrids (Figure 1A). Furthermore, accumulation of these lipids is non-linear, and changes most abruptly at the transition between encased and emerged silks (i.e., section C to D) and along the emerged portions of silks (i.e., section D to E) (Figure 1A). Indeed, regression of the cuticular lipid accumulation along the silk length reveals that a quadratic polynomial model (adjusted R^2^ ranging from 0.2 to 0.9) is a better fit to the dynamics of total lipids, hydrocarbons, VLCFAs, and aldehydes as compared to a linear model (Supplemental Table S2).

The compositional dynamics of these chemical classes vary not only along the silk length, but also among the different genotypes. Silks from inbred B73 and hybrid B73×Mo17 exhibited a decrease in the relative abundance of VLCFAs and a concomitant increase in relative abundance of hydrocarbons along the silk length (Figure 1B). In contrast, there was no change in Mo17×B73 and no consistent change for Mo17 in relative abundance of VLCFAs along the silk (Figure 1B), but for each silk section, cuticular VLCFA concentration for Mo17 and both reciprocal hybrids was 1.7-to 4.7-fold higher than B73. The largest difference among the genotypes occurs for emerged silks (sections D and E) (Figure 1A), suggesting that VLCFAs in Mo17 are preferentially transported to the silk surface instead of being utilized for hydrocarbon production.

Hydrocarbon composition, specifically the degree of unsaturation, also varies among these genotypes. Although alkenes consistently comprise ∼50% of the hydrocarbons along the spatio-temporal gradient of the silks for Mo17 and the reciprocal hybrids, for B73 the relative abundance of alkenes increased from approximately 40% at the base to 50% of the hydrocarbons at the tips of the silks (Figure 1B). Most alkenes harbor a single double bond (i.e. a monoene), which is positioned either at the 7^th^ or 9^th^ carbon-position, irrespective of the alkyl chain-length (Figure 1B). However, additional minor monoenes were identified with double bonds positioned at the 14^th^ or 15^th^ carbon-positions, within hydrocarbons of 29, 31, and 33 carbon atom chain lengths. Accumulation patterns of the 7- and 9-monoenes differed between the inbred lines (Figure 1B). Specifically, for B73, 7-monoenes consistently comprise ∼50% of all alkenes along the spatio-temporal gradient, while the relative abundance of 9-monoenes increased from 33% to 42% (Figure 1B) of the alkenes along this gradient. In contrast, on Mo17 silks, the relative abundance of both the 7- and 9-monoenes did not change along the spatio-temporal gradient, and they accounted for 64% and 20% of all alkenes, respectively (Figure 1B). Thus, although the absolute amounts (µmol/g dry weight tissue) of these alkenes increased by ∼2.5-fold in the spatio-temporal gradient of the silks (Supplemental Figure S1), the relative proportions of 7- and 9-monenes remained constant for Mo17, with the 9-monoenes being less abundant (Figure 1B).

The relative impacts of genotype, silk development and silk encasement status (i.e. silk microenvironment) on the dynamics of the cuticular lipid metabolome were visualized via Principal Component Analysis (PCA) of the 45 cuticular lipid metabolites (Figure 1C). The distribution of silk samples along PC1, which accounts for 31% of the total variance, is tightly associated with the position along the length of silks, and mainly separates the husk-encased (sections A-C) from the emerged (sections D and E) portions of the silks. Moreover, PC1 also separates based upon silk development, separating section C from sections A and B within the husk-encased silks of the inbred B73 and the two hybrids, as well as separating sections D and E of the emerged silks from all evaluated genotypes. PC2 accounts for 19% of the total variance and primarily separates genotypes, with inbred lines B73 and Mo17 forming subclusters, and the hybrids forming a third cluster, which is closer to Mo17 than B73.

Hence, we hypothesize that the spatio-temporal gradient of the silk has a greater influence on cuticular lipid concentrations than the genetic variation that was evaluated in this study. Two-way ANOVA on specific lipid classes (i.e. hydrocarbon, aldehyde and alcohol) and each individual metabolite (Supplemental Table S3; Supplemental Table S4) supports this hypothesis and reveals that the position along the length of the silk is the major determinant of the observed variance in total cuticular lipid accumulation (accounting for 70% of the variance). The effect of genotype and the two-way interaction between genotype and silk section explain a much smaller proportion (between 5% and 10%) of the variance in the accumulation of these components (Supplemental Table S4). In stark contrast, but in agreement with the major patterns in Figure 1B, the majority of the observed variance in cuticular VLCFA accumulation is explained primarily by genotype (66%), whereas the position along the length of the silk explains only 17% of this variation.

### Silk microenvironment is the major driver of changes in cuticular lipid accumulation

The variation in the cuticular lipid metabolome along the silk length may be explained by either or both the silk developmental gradient and/or the change in the microenvironment of the silks associated with husk-encasement status. At the time of silk collection, cell division has ceased, and the developmental gradient is primarily determined by cellular elongation, which is occurring only in the husk-encased portions of the silk, with an acropetal decrease in the rate of cell elongation (Fuad-Hassan et al., 2008). The potentially confounding determinants of silk cellular development and silk encasement status were untangled by the application of a nested ANOVA statistical test. Silk encasement status explains 57% of the variation in total cuticular lipid abundance, whereas the silk acropetal gradient (i.e., sections A *versus* B *versus* C for encased portions, and section D *versus* E for emerged portions) explained only 13% of the observed variance (Supplemental Table S4). One can conclude therefore, that cuticular lipid accumulation is primarily impacted by the change in silk microenvironment associated with emergence of the silks from the protective husk leaves.

### Correlations among cuticular lipid metabolites along the silk length reveal genotype-dependent clustering patterns

Pairwise Spearman correlation matrices between each lipid metabolite pair along the length of silks were subjected to weighted gene correlation network analysis (WGCNA) (Horvath and Langfelder, 2011). WGCNA is a method that was first developed for identifying clusters of co-expressed genes, but is also now widely applied to metabolomics data (Shen et al., 2013; Bartzis et al., 2017; Rosato et al., 2018). The cluster membership of many metabolites differs among the four genotypes that were evaluated (Figure 2; Supplemental Figure S2). In the inbred B73, and the reciprocal hybrids, the majority of metabolites group into a single cluster (Table 1; Figure 2A; Supplemental Figure S2) that includes hydrocarbon products (*HC_2n-1_* and *HC_2n_*) and corresponding precursor VLCFAs (*FA_2n_* and *FA_2n+1_*). In contrast, in the inbred Mo17, hydrocarbons and VLCFAs mostly reside in separate clusters (Table 1; Figure 2, B and C). These observed differences in correlation network topologies suggest that the dynamics of cuticular lipid accumulation differ between Mo17 and B73, especially the dynamics of VLCFA accumulation.

**Figure 2.**
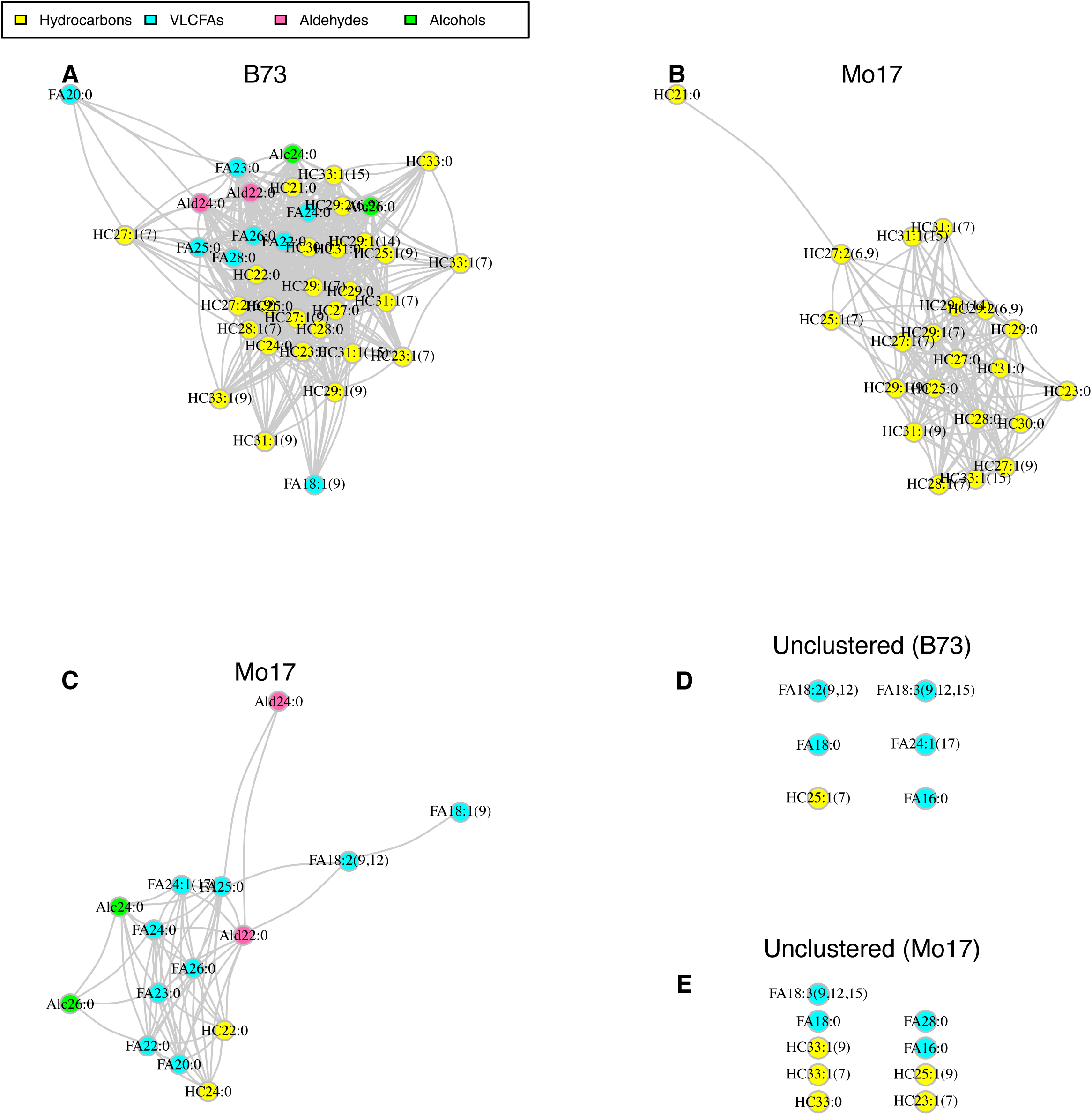
Correlation-based clustering of silk cuticular lipid abundance data for inbred lines, B73 and Mo17. Rank-based Spearman correlations were calculated between all pairs of metabolites and used to construct the weighted correlation networks via WGCNA for **A**, B73 and **BC**, Mo17. The non-clustered metabolites for B73 and Mo17 are shown in **D** and **E**, respectively. Pairs of lipid metabolites connected by edges are significantly correlated with correlation coefficients ≥0.5 and reside within the same cluster. Edge length represents correlation strength with shorter edges representing stronger correlations between metabolites. Unclustered singleton metabolites were not statistically correlated with any other metabolites, or shared correlation values <0.5.

**Table 1.**
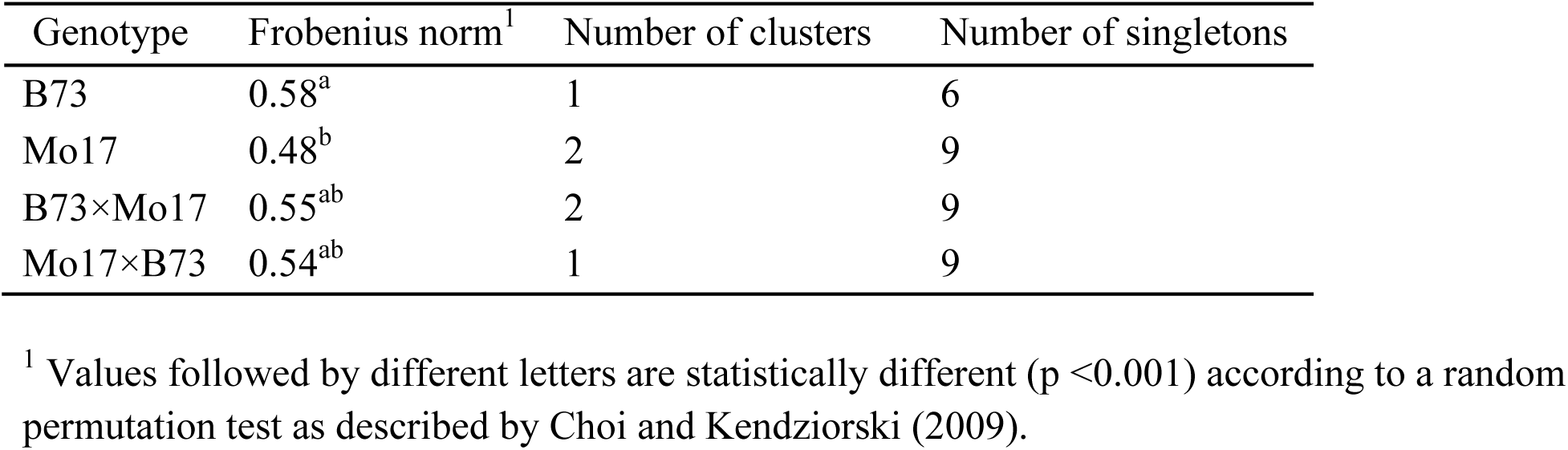
Silk cuticular lipid metabolites from B73 and Mo17 exhibit different correlation patterns along the spatio-temporal silk gradient. The Frobenius norm is derived from the sum of squares for all the elements in a Spearman correlation matrix. Correlation-based clustering was performed by WGCNA.

| Genotype | Frobenius norm<sup>1</sup> | Number of clusters | Number of singletons |
| --- | --- | --- | --- |
| B73 | 0.58<sup>a</sup> | 1 | 6 |
| Mo17 | 0.48<sup>b</sup> | 2 | 9 |
| B73×Mo17 | 0.55<sup>ab</sup> | 2 | 9 |
| Mo17×B73 | 0.54<sup>ab</sup> | 1 | 9 |

<sup>1</sup> Values followed by different letters are statistically different (p <0.001) according to a random permutation test as described by Choi and Kendzierski (2009).

This conclusion is confirmed by comparison of the Frobenius norms of the Spearman correlation matrices, which are aggregate measures of the level of correlation among all metabolite pairs as they are modulated by the spatio-temporal silk gradient of a given genotype. The Frobenius norms for silks of B73 and the reciprocal hybrids were statistically higher than for silks of Mo17 (Table 1), demonstrating that there are stronger correlations among the profiled metabolites in each of the former genotypes as compared to Mo17.

A commonality among all genotypes, however, is the lack of correlation of 16- and 18-carbon FAs with the majority of the cuticular lipid metabolites. Notably, these FAs are found either as singleton, unclustered metabolites (B73, Mo17, Mo17×B73) or within a single small cluster (B73×Mo17) (Figure 2, D and E; Supplemental Figure S2), suggesting that the metabolism of these fatty acids is distinct from the other cuticular lipid metabolites, i.e. hydrocarbons and VLCFAs with ≥21 carbons.

### Identification of the signature metabolites that are primary contributors to compositional variation in the cuticular lipid metabolome

Signature cuticular lipid metabolites (or biomarkers) were identified via multivariate Partial Least Squares-Discriminant Analysis (PLS-DA). This statistical strategy is commonly applied to metabolite data for classification analysis and biomarker selection (Worley and Powers, 2012; Bartel et al., 2013), and identified the signature metabolites that most contribute to the variation in the cuticular lipid metabolome as modulated by the change in the silk microenvironment or by the silk genotype. In this study, two supervised PLS-DA regression models were constructed, with the explanatory variables (R^2^X) being the concentrations of individual cuticular lipid metabolites and the response variable (R^2^Y) being either genotype (Supplemental Figure S3, A and C) or silk section (Supplemental Figure S3, B and D). The corresponding weight plots, illustrating the contributions of each metabolite in discriminating among genotypes or silk sections, are presented in Supplemental Figure S3, C and D, respectively. The metabolites that contribute the most to PLS-DA classification were identified via a variable-importance-in-projection score (VIP>1; Supplemental Table S5), which is a cumulative measure of the contribution of a given explanatory variable (i.e., metabolite) to a PLS-DA model (Pérez-Enciso and Tenenhaus, 2003).

The PLS-DA classification of metabolome compositions relative to genotype yields two clusters of silk samples for each of the inbred lines, and a third cluster containing the samples from the two hybrids (Supplemental Figure S3A). The resulting model explained 63% and 66% of the variance in the explanatory (R^2^X) and response (R^2^Y) variables, respectively. The predictive accuracy is relatively low for this model (i.e., Q^2^Y = 63%), due to the inability to correctly classify the metabolite compositions between the two hybrids, B73×Mo17 and Mo17×B73, which express similar cuticular lipid metabolomes. However, upon combining the metabolite data from the two hybrids into a single group, the predictive accuracy of the model was improved (Q^2^Y = 88%). As visualized in the PLS-DA scatter plot, the signature lipids (i.e., VIP scores >1) identified in the original PLS-DA model are major contributors to sample classification, primarily differentiating B73 from Mo17, and these two parental lines from the B73×Mo17 and Mo17×B73 hybrids (Supplemental Figure S3A; black circles). These signature metabolites (10 fatty acids and 5 hydrocarbons) (Figure 3, A and C) are one third of the total number of detected cuticular lipid metabolites but account for two thirds of the observed variance in the response variable.

**Figure 3.**
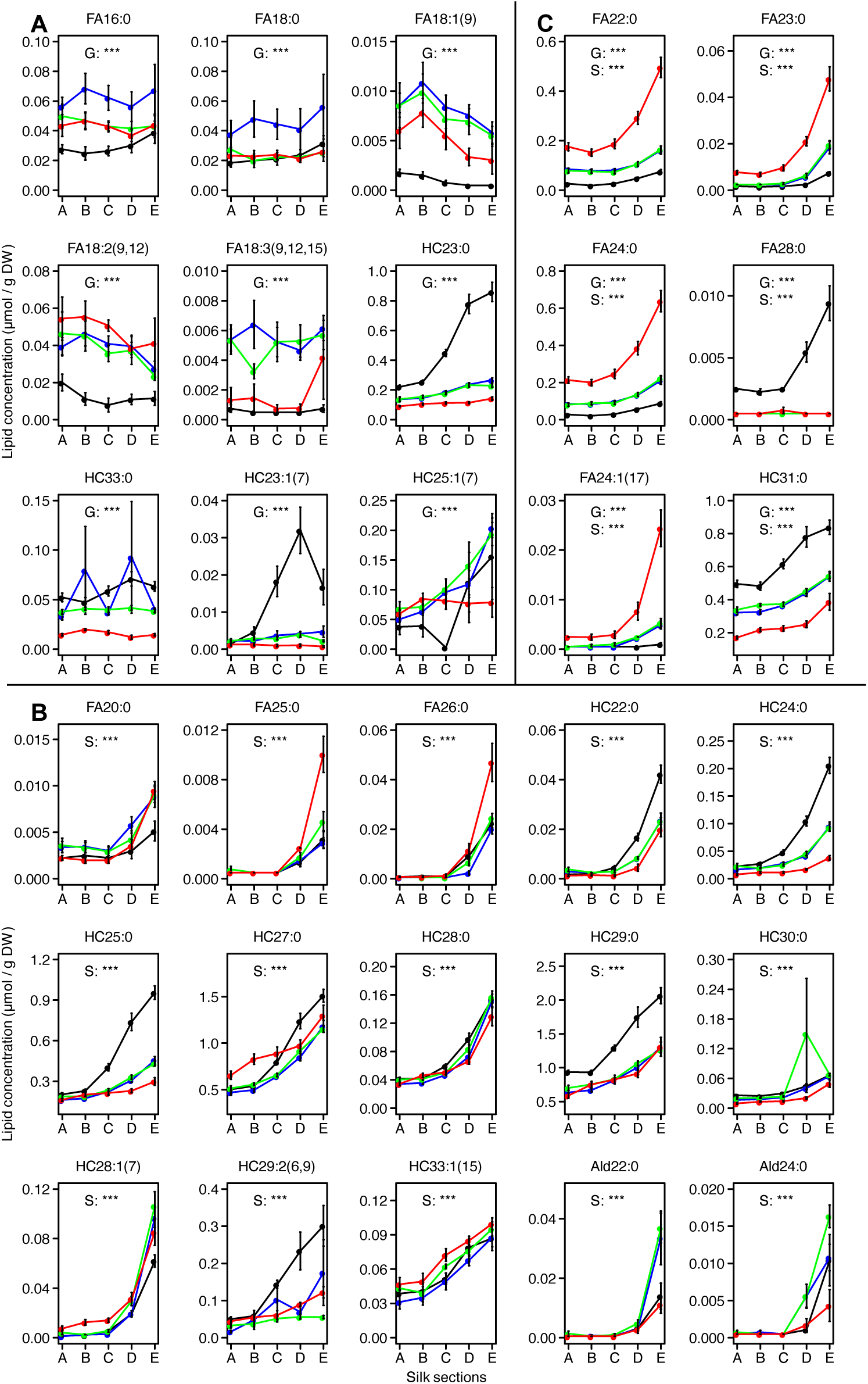
Accumulation patterns of signature cuticular lipid metabolites that distinguish among genotypes or among silk sections. Concentrations of signature cuticular lipids identified by partial least square discriminant analysis (PLS-DA) as having variable-importance-in-projection (VIP) scores >1. Nine (**A**) and fifteen (**B**) of these metabolites contribute to either the observed genotype-based or silk section-based PLS-DA separation, respectively, and an additional six lipid metabolites were selected in both categories (**C**). Two-way ANOVA of the main effects of genotype and silk section were conducted for each metabolite, and statistical significance is noted with asterisks (***, p<0.0001) for genotype (G), silk section (S) or both G and S. The interaction effect, G × S was also evaluated by ANOVA for each metabolite and presented in Supplementary Table S3. Averages ± SE are reported, with averages connected by colored lines to facilitate visualization.

The PLS-DA classification model of metabolome compositions relative to the spatio-temporal silk gradient yields one cluster that is specific to husk-encased sections A and B, and three individual clusters specific to silk sections C, D, and E (R^2^X=55%, R^2^Y=31%, Q^2^Y =28%; Supplemental Figure S3B). A PLS-DA model that distinguishes silk sections based on husk-encasement status improves the predictive accuracy (Q^2^Y) from 28% to 79%, demonstrating the importance of the silk microenvironment in shaping the cuticular lipid metabolome. Moreover, the PLS-DA model that classifies the metabolomes by silk sections, identified 21 signature metabolites (8 fatty acids, 2 aldehydes and 11 hydrocarbons) (Supplemental Figure S3D, black circles), each of which exhibited statistically significant differences in patterns of accumulation among the silk sections (Figure 3, B and C). In combination, these signature metabolites account for 75% of the variance observed for silk sections. Notably, six metabolites (four saturated fatty acids, an unsaturated fatty acid, and an alkane) were selected as signature metabolites that contribute to the variation observed across both different genotypes and different silk sections (Figure 3B). Collectively, these quantitative statistical analyses demonstrate the importance of both the silk microenvironment and the silk genotype on determining the cuticular lipid composition, and further identify that the accumulation of many of the VLCFA precursors, aldehyde intermediates and hydrocarbon products are differentially influenced by these two factors.

### Cellular free VLCFAs and cuticular VLCFAs are positively correlated

The VLCFAs that contribute to the assembly of the silk cuticle are products of the endoplasmic reticulum-associated fatty acid elongase, and these VLCFAs can also be utilized to assemble other cellular complex lipids (e.g., phospholipids, neutral lipids and ceramide lipids). Therefore, we sought to explore the relationship between cellular VLCFAs and those associated with cuticular lipids. Hence, in parallel to the preparation of cuticular lipid extracts from silks of B73 and Mo17 inbreds (Figure 4A), total cellular lipids were extracted and analyzed from these silks (Figure 4, B and C). The cellular lipids were profiled by liquid chromatography-mass spectrometry (LC-MS), which identified the occurrence of free VLCFAs and VLCFAs that were associated with complex lipids. The cellular complex lipid VLCFAs were associated with three ceramides, containing FA_22:0_, FA_24:0_, or FA_26:0_, one phosphatidylcholine molecular species that contained FA_20:0_ and one triacylglycerol molecular species that contained FA_22:0_. The free VLCFAs included eight saturated VLCFAs of 20-to 34-carbon chain lengths (Supplemental Table S6).

**Figure 4.**
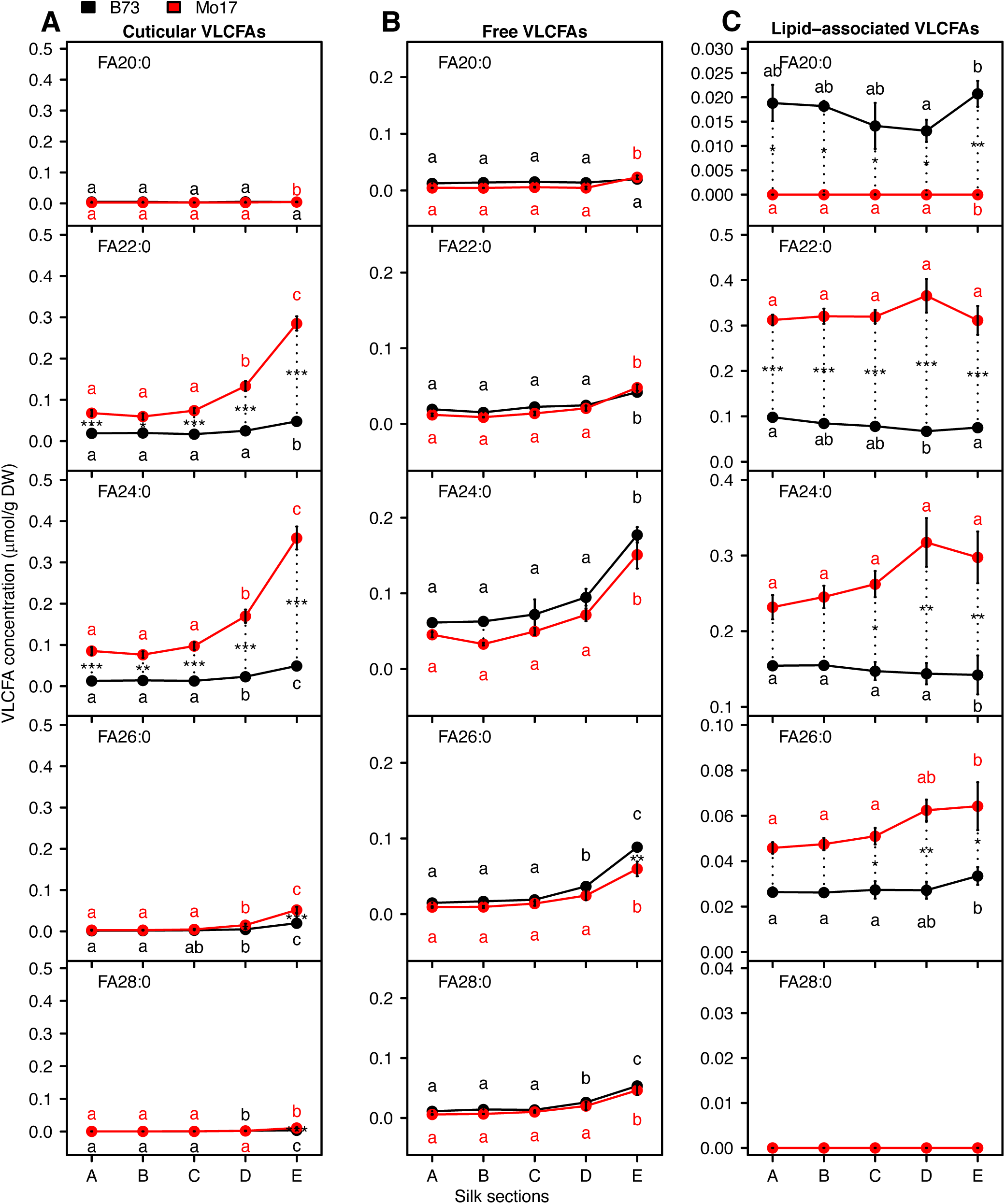
Accumulation of three VLCFA pools along the silk spatio-temporal gradient. Concentrations of individual cuticular VLCFAs (**A**), cellular free VLCFAs (**B**) and cellular lipid-associated VLCFAs (**C**) in different silk sections of inbred lines B73 (black data points) and Mo17 (red data points). For each genotype, different letters associated with data points indicate statistical differences among silk sections (p <0.05; Tukey’s HSD). Asterisks denote statistical differences between genotypes at a specific position along the silk length (***, p <0.0001; **, p <0.001; *, p <0.05). Averages ± SE from three to four replicates are reported for the VLCFA pools; N=35.

Of the three VLCFA pools that were evaluated (i.e., cellular free VLCFAs, lipid-associated VLCFAs, and cuticular VLCFAs), the cellular free VLCFA fraction increased in abundance from the base to the tip of the silks (Figure 4). This increase in cellular free VLCFAs correlated with the cuticular VLCFA fraction on silks of both B73 (Pearson correlation coefficients, 0.65 to 0.92) and Mo17 (0.87 to 0.94), whereas no such correlation was observed with the cellular lipid-associated VLCFAs for either genotype (Supplemental Table S7). These results are consistent with the model that the cellular free VLCFA pool contributes to the accumulation of VLCFAs in the silk cuticle.

### Cuticular lipid product-precursor ratios vary among genotypes and across acyl chain lengths

The dynamics of the hydrocarbon-producing pathway were explored by comparing the steady state abundances of cuticular hydrocarbon products and the corresponding VLCFA precursors that accumulate on the silk surface. The signature cuticular lipids that distinguish either or both genotypes and silk sections include six pairs of saturated or unsaturated hydrocarbons and fatty acids (i.e., either *HC_(2n-1):0_* and *FA_2n:0_,* or *HC_(2n-1):1_* and *FA_2n:1_* metabolites) (Supplemental Figure S4). These can be considered “product-precursor” pairs of parallel hydrocarbon producing pathways (Perera et al., 2010). There is a striking difference in the accumulation of these VLCFAs as compared to the hydrocarbons along the silk spatio-temporal gradient in the two inbreds. Namely, as compared to B73 silks, at the more prevalent carbon chain lengths, VLCFAs are 2-to 13-fold more abundant in Mo17, whereas hydrocarbons are 2-to 13-fold less abundant in Mo17 (Supplemental Figure S4). These distinct accumulation patterns of hydrocarbons and VLCFAs between the two inbreds suggest that the conversion of each VLCFA to the corresponding hydrocarbon may not proceed to the same extent in Mo17 silks, as compared to B73 silks.

Further interrogation of the product-precursor ratios revealed that precursor chain length was the most influential factor on these ratios. In the four genotypes examined, the saturated VLCFA (*FA_2n:0_*) of each product-precursor pair decreased in abundance with increasing carbon chain length, concomitant with an increase in the corresponding alkane product (*HC_2n-1:0_*) (Figure 5; Supplemental Figure S5). Indeed, regression analysis of the product-precursor ratio (i.e., *HC_2n-1:_*_0_ : *FA_2n:0_*) confirms a linear increase in the value of this ratio as the carbon chain length increases from 22 to 28 (p-values <0.0001, R^2^≥0.79; Figure 5; Supplemental Figure S5), and this is independent of silk husk-encasement status or genotype. This statistically significant linear relationship suggests that with increasing acyl chain length, silk VLCFAs are increasingly recruited as precursors for the biosynthesis of hydrocarbon products.

**Figure 5.**
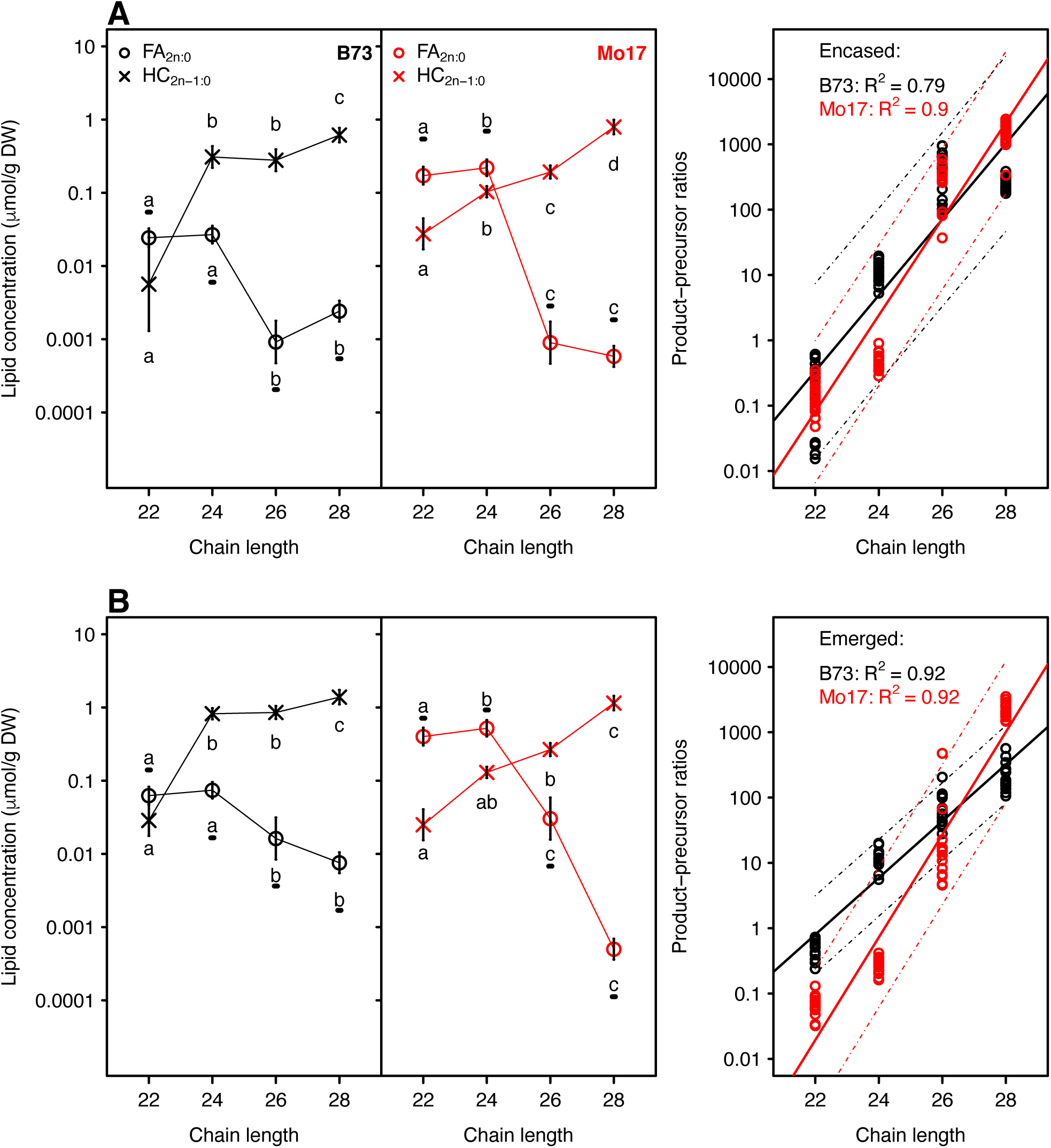
Accumulation of cuticular hydrocarbon products and corresponding cuticular VLCFAs, and the regression of the product-precursor ratios on chain length for inbreds B73 and Mo17. Concentrations (log-scaled) of cuticular hydrocarbons (*HC_2n-1:0_*) and VLCFAs (*FA_2n:0_* ) were analyzed from husk-encased silks (**A**) and from silks that had emerged from the husks (**B**) of the inbreds B73 (black data points) and Mo17 (red data points). Averages ± SE from seven to eight replicates are reported for the metabolite concentrations. Different letters associated with data points from the same metabolite class indicate a statistically significant difference between acyl chain lengths (p <0.05; Tukey’s HSD test); letters associated with VLCFAs are underlined. In the regression, the prediction intervals for the regression models are indicated by the dashed lines for each inbred.

Based on the observations that this product-precursor ratio increases with increasing chain length (Figure 5) and that cuticular VLCFAs of longer chain lengths are less abundant on the silk surface (Supplemental Table S1), Bayesian model selection was used to interrogate the other factors that may affect this ratio. Specifically, while keeping metabolite chain length (*2n*) in the baseline model, our goal is to establish the relative importance of the other variables (i.e., genotype, husk-encasement status, growing year, cellular free VLCFAs, and lipid-associated VLCFAs on the product-precursor ratio). The construction of Bayesian regression models and the methods of their comparisons are detailed in Supplementary Methods. Because we did not have a single dataset that integrates the factors of both cellular VLCFAs and growing year, we performed two Bayesian Model Comparisons, examining the impact of cellular VLCFAs on the product-precursor ratio (i.e., VLCFA model selection) or the impact of growing year on this ratio (i.e., Environment model selection).

To assess the impact of cellular VLCFAs on product-precursor ratios, we considered three Bayesian regression models: VLCFA model 1, which includes cellular free-VLCFAs; VLCFA model 2, which includes cellular lipid-associated VLCFAs; and VLCFA model 3, which includes both cellular free and lipid-associated VLCFAs (Table 2). The predictor variables included in each VLCFA model were selected by preliminary Bayesian Model Comparison (see Supplemental Methods; Supplemental Table S8) from the variables genotype, husk-encasement status, and metabolite chain length, as well as those associated with two-way and three-way interaction terms among these predictor variables. Subsequently, a second round of Bayesian Model Comparison was performed to compare a model without VLCFA variables (Baseline Model) to VLCFA models 1, 2, and 3. The resultant Bayes factors demonstrate a substantial increase in the ability of each of the VLCFA models to explain patterns of product-precursor relationships when incorporating cellular VLCFAs, be they free or lipid-associated VLCFAs (Table 2). However, the incorporation of free VLCFAs in the model (i.e., VLCFA model 1) results in the highest Bayes factor (1.88×10^7^), demonstrating that the cellular free VLCFAs have the highest influence on the product-precursor ratios of the cuticular lipids. Furthermore, the high correlation between the cellular free VLCFAs and the cuticular VLCFA pool (Supplementary Table S7) is consistent with the precursor role of very long chain fatty acyl-CoAs as the precursors for cuticular hydrocarbon biosynthesis.

**Table 2.** Impacts of cellular free VLCFAs and lipid-associated VLCFAs on product-precursor ratios determined by Bayesian model comparison. The product-precursor relationship was represented by the linear regression of product-precursor ratio (*HC_2n-1_* : *FA_2n_*) *versus* acyl chain length, *2n.* Bayesian model comparison was performed on metabolomics data collected during the 2015 growing year with cellular free VLCFAs and cellular lipid-associated VLCFAs profiled from lipids extracted from the entire silk tissue. The notation for cuticular lipids is described in Methods.

| Baseline model | VLCFA models<sup>a</sup> | Examined terms | Bayes factor<sup>b</sup> |
| --- | --- | --- | --- |
| Genotype ($g$) + Encasement status ($k$) + Acyl chain length ($n$) + ($g \times k$) + ($k \times n$) + ($g \times k \times n$) | *VLCFA model 1:* $g + k + n + (g \times k) + (k \times n) + (g \times k \times n) + [\text{free VLCFAs } (ffa) + (ffa \times g) + (ffa \times k) + (ffa \times n) + (ffa \times k \times n)]$ | Cellular free VLCFAs ($ffa$) and the interaction effects with $g$, $k$ and $n$ | $1.88 \times 10^7$ |
| | *VLCFA model 2:* $g + k + n + (g \times k) + (k \times n) + (g \times k \times n) + [\text{lipid-associated VLCFAs } (lfa) + (lfa \times g) + (lfa \times n)]$ | Cellular lipid-associated VLCFAs ($lfa$) and the interaction effects with $g$ and $n$ | $5.31 \times 10^6$ |
| | *VLCFA model 3:* $g + k + n + (g \times k) + (k \times n) + (g \times k \times n) + [ffa + (ffa \times g) + (ffa \times k) + (ffa \times n) + (ffa \times k \times n) + lfa + (lfa \times g) + (lfa \times n)]$ | Cellular free VLCFAs ($ffa$) and lipid-associated VLCFAs ($lfa$), and interaction effects with $g$, $k$ and $n$ | $4.57 \times 10^6$ |

<sup>a</sup> For each VLCFA model, the main effect of the examined term and the associated interaction effects were added to the baseline model as justified in Supplementary Table S8.
<sup>b</sup> Bayes factors were calculated as the ratio of the likelihood between VLCFA models (numerator) and the baseline model (denominator). A Bayes factor $>3.2$ indicates that the addition of the examined terms to the baseline model resulted in significantly better fit to the data.

Subsequently, using combined data collected from the two growing years, we queried the effect of growing year on product-precursor relationships in the context of the different genotypes and silk encasement status (Table 3, Environment model selection). Because the cellular VLCFA pools were not measured in the first growing year, free VLCFAs, lipid-associated VLCFAs and the related interaction terms were not included in this modeling effort. The genotype factor exhibited the strongest impact on this product-precursor relationship in the combined dataset (Bayes factor = 1.01×10^110^, Table 3, Environment model selection). Growing year (Bayes factor =2.35×10^33^) and husk-encasement status (Bayes factor = 7.76×10^28^) also impacted the product-precursor ratios, although to a lesser extent. Similarly, such quantitative modeling analyses indicate that the two-way interactions between genotype and husk encasement status, and between genotype and growing year also play a role in determining the product-precursor ratio dynamics along the silk spatio-temporal gradient (Supplemental Table S2), but husk encasement status and year does not. Collectively therefore, these analyses illustrate the complexity of factors that influence product-precursor relationships with respect to alkyl carbon chain length of cuticular lipid components.

**Table 3.**
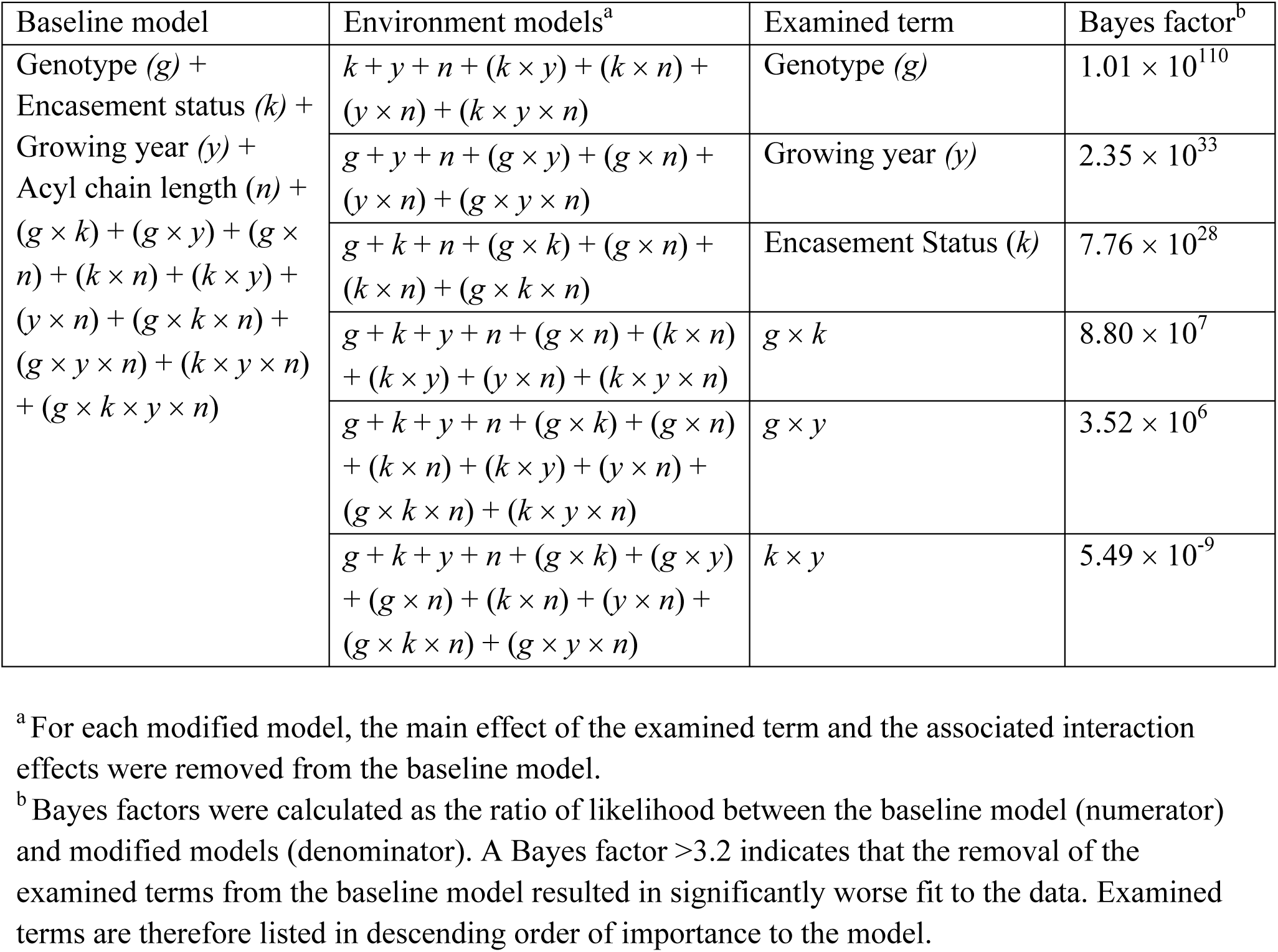
Impact of genotype, husk-encasement status, and growing year on the product-precursor ratios determined by Bayesian model comparison. The product-precursor relationship was represented by the linear regression of product-precursor ratio (*HC_2n-1_* : *FA_2n_*) *versus* the alkyl chain length, *2n*. Bayesian model comparison was performed on the combined metabolomics datasets gathered from tissue grown in 2014 and 2015.

## DISCUSSION

In this study we utilized cuticular lipid profiling data gathered from the spatio-temporal gradient of maize silks to assess and compare the dynamics of the metabolic network that supports cuticular hydrocarbon accumulation. Previously, a series of parallel pathways has been proposed for the biosynthesis of these hydrocarbons, which are primarily composed of alkanes and alkenes (Perera et al., 2010). These pathways involve elongation of fatty acids, followed by reduction and decarbonylation to generate a homologous series of alkanes. In parallel, a homologous series of alkenes can be generated from the reduction and decarbonylation of unsaturated fatty acids that either undergo desaturation followed by elongation, or elongation followed by desaturation.

We investigated the cuticular lipid metabolomes for the maize B73 and Mo17 inbred lines, and the corresponding reciprocal hybrids. Genotypic variation of cuticular lipid compositions occurs primarily between the two inbred parents (i.e., B73 *versus* Mo17) and between the inbred parents and the hybrids, but not between the reciprocal hybrids, B73×Mo17 and Mo17×B73. While some plant molecular phenotypes (e.g., chloroplast thylakoid lipid composition and localization; Dueñas et al., 2017) and gene expression profiles (Gonzalo et al., 2007; Swanson-Wagner et al., 2009) can differentiate these reciprocal hybrids, such observations are less common than the differences that occur between the parental inbred lines (Paschold et al., 2012; Baldauf et al., 2016; Marcon et al., 2017). The remainder of the discussion is therefore focused on the cuticular lipid metabolomes and the apparent variation in the underlying hydrocarbon-producing pathway between the two inbred parents.

### Maize silks as a novel system to study the dynamics of the cuticular lipid biosynthesis pathway

Maize silks provide a unique platform for assessing the dynamics of hydrocarbon production, in the absence of complex flux analysis approaches. Many characterizations of the cuticular lipid biosynthesis pathway have been conducted for single time-point comparisons of metabolomes profiled from different genetic backgrounds (as reviewed by Xue et al., 2017 and Trivedi et al., 2019). In this study, cuticular lipid profiling captures differences in accumulation of precursors, intermediates, and products of the pathway along a spatio-temporal gradient that includes the developmental progression of silks (Fuad-Hassan et al., 2008) as well as an environmental transition as portions of the silks change encasement status from husk-encased to husk-emerged, when silks become exposed to the external environment. Similar spatio-temporal studies have identified metabolic shifts in primary, secondary, and lipid metabolism along the lengths of leaves in Arabidopsis, tobacco, wheat and maize (Pick et al., 2011; Watanabe et al., 2013; Allwood et al., 2015; Li et al., 2017; Zhou et al., 2019; Bourgault et al., 2020).

In the case of maize silks, the hydrocarbon end-products of cuticular lipid biosynthesis accumulate extracellularly on the silk surface (Perera et al., 2010; Loneman et al., 2017), whereas VLCFAs accumulate both internally as either free VLCFAs or lipid-associated VLCFAs (e.g. triacylglycerols and ceramides), and on the silk surface as components of the cuticle. Here we show that the concentrations of individual fatty acids that accumulate on the silk surface are highly correlated with the concentrations of corresponding free cellular VLCFAs. The exceptions to this observation are the 18-carbon fatty acids (i.e. stearic and oleic acids), whose concentrations are poorly correlated between the cellular lipid fraction and the cuticular lipid fraction. This lack of correlation is attributable to the role of 18-carbon fatty acids as initial precursors for elongation of VLCFAs (Bach and Faure, 2010), and as building blocks in the assembly of numerous complex lipids, including for membrane glycerolipids (Sato and Awai, 2017), phospholipids (Nakamura, 2017), and sphingolipids (Michaelson et al., 2016). Indeed, an appreciable amount of saturated and unsaturated 18-carbon fatty acids profiled from silk tissue were associated with complex cellular lipids, such as triacylglycerol or phosphatidylcholine. In contrast, VLCFAs are components of cuticular lipids, the ceramide component of sphingolipids (Dietrich et al., 2005) and some discrete phospholipids (Michaelson et al., 2016), which is consistent with our observation that VLCFAs are recovered from the silk cuticle and as components of three specific cellular glucosylceramide species harboring a single hydroxylated C22-, C24-or C26-VLCFA moiety. Collectively, these observations suggest that the spatio-temporal accumulation of VLCFAs on maize silks reflects the dynamics of the steady-state levels of the cellular free VLCFA precursor pool utilized by the hydrocarbon-producing pathway.

### The silk microenvironment significantly impacts the cuticular lipid metabolome and the hydrocarbon-producing pathway

Emergence of the silks from the encasing husk-leaves has previously been shown to induce increased accumulation of cuticular hydrocarbons in different maize genotypes (Yang et al., 1992; Miller et al., 2003; Perera et al., 2010; Loneman et al., 2017; Dennison et al., 2019). However, in each of these prior studies that compared the cuticular lipid metabolome between husk-encased and emerged portions of silks, the change in microenvironment (i.e. silk encasement status) was overlaid with a confounding developmental gradient, particularly that of an acropetal decrease in cellular elongation within the husk-encased silks at the time of tissue sampling (Fuad-Hassan et al., 2008). This study shows that the change in microenvironment *per se* (i.e. husk-encased vs emerged silks) drives the observed change in cuticular lipid composition, as well as the product-precursor relationships within the cuticular hydrocarbon-producing pathway. Indeed, a recent transcriptomic atlas for maize silks has identified microenvironment-based differential gene expression for many genes involved in cuticular lipid biosynthesis, and more broadly for genes involved in responses to biotic and abiotic stresses (McNinch et al., 2020). Interestingly, the increase in cuticular hydrocarbon accumulation that is observed in emerged silks of B73, the B73×Mo17 and Mo17×B73 hybrids and, to a lesser extent, Mo17, may provide protection against drought, as for example, the observed increase in alkane accumulation upon drought stress that occurs in soybean and sesame (Kim et al., 2007a; Kim et al., 2007b).

### Supervised and unsupervised multivariate analyses identified key factors that impact the hydrocarbon-producing pathway

In metabolomic studies, correlation networks are often used to identify metabolites that are co-regulated in response to perturbations caused by disease, chemical and other environmental treatments, and genetic modifications (DiLeo et al., 2011; Fukushima et al., 2011; Papazian et al., 2016; Angelovici et al., 2017). Such correlation networks have been constructed from non-targeted metabolomics data gathered from different maize inbred lines, which capitulate pathways of primary metabolism in leaves (Toubiana et al., 2016; Zhou et al., 2019) and kernels (Rao et al., 2014). However, the correlative relationships of metabolites within targeted metabolomes are more complex, as queried metabolites are often from highly related pathways, and therefore the accumulation patterns of most metabolites are inherently correlated. Even so, such correlation networks have successfully described a change in metabolic states due to change in development-, genotype-, or stress-specific impacts (Steuer, 2006; Sawada et al., 2009; Warth et al., 2015; Li et al., 2016; French et al., 2018). In our correlation analyses of metabolites involved in the hydrocarbon-producing network, hydrocarbon products and corresponding precursor VLCFAs separated into different clusters within the correlation network in Mo17, whereas these products and precursors are highly correlated in B73.

The combination of metabolite profiling and subsequent statistical analysis allows for the identification of signature metabolite biomarkers that represent changes in metabolic status induced by genetic or environmental perturbations, or by tissue development (Fernandez et al., 2016; Hong et al., 2016). For example, metabolite biomarkers have successfully uncovered variations within complex pathways in response to developmental or environmental stimuli in Arabidopsis (Sulpice et al., 2009), rice (Tarpley et al., 2005), maize (Obata et al., 2015; Luo et al., 2019), *Brassica nigra* (Papazian et al., 2016) and wheat (Kang et al., 2019). Herein, we identified 30 cuticular lipids as signature metabolites that include six pairs of cuticular VLCFAs and hydrocarbon products. The spatio-temporal accumulation pattern of these signature metabolites demonstrates that the cuticular lipid-producing network is enhanced along the spatio-temporal gradient of the silks, especially during the transition of the microenvironment experienced by the silk tissue caused by emergence from the encasing husks. Furthermore, a comparison of the concentrations of the signature metabolites between B73 and Mo17 suggest that the genetic difference of the silks is key in determining the product-precursor relationship of the cuticular lipid-producing network. This statistical approach for analyzing metabolomics data that combined supervised and unsupervised multivariate analyses not only provides new insights into the factors that impact cuticular lipid accumulation and composition, but also permits probing the status of the individual metabolite conversions within the hydrocarbon-producing pathway.

### Factors impacting the product-precursor relationship

Product-precursor relationships have previously been inferred from cuticular lipid profiles of several systems. These include the relationships between alkanes, secondary alcohols, and ketones, and the relationships between VLCFAs, alcohols and wax esters of Arabidopsis cuticles (Lai et al., 2007; Wen and Jetter, 2009), the relationships between primary and secondary diols and esters of the wheat cuticle (Racovita and Jetter, 2016), and relationships between unsaturated aldehydes and alkenes of the maize cuticle (Perera et al., 2010). In each case, the presence of precursors that are chemically related by a potential biochemical reaction was posited to demonstrate biosynthetic relationships amongst these metabolites.

In this study, we compared the accumulation pattern of cuticular alkanes (*HC_2n-1:0_*), and the associated VLCFAs (*FA_2n:0_*) that can serve either as precursors in the pathway or can themselves be deposited as cuticular lipid products. More specifically, we considered that the individual VLCFAs queried in this study can serve as precursors for three processes: (1) fatty acid elongation that generates longer chain VLCFAs, (2) sequential reduction-decarbonylation reactions that generate alkanes, or (3) export to the silk cuticle to generate the cuticular VLCFA pool. Along the spatio-temporal gradient of the silks, both cuticular and total cellular VLCFAs increase in accumulation. Thus, VLCFA biosynthesis is likely promoted in the distal end of the spatio-temporal gradient of silks, resulting in increased cuticular lipid load as either free VLCFAs or VLCFA-derived hydrocarbons produced by a series of elongation-reduction-decarbonylation reactions (Perera et al., 2010).

A Bayesian approach has previously been employed to predict protein trafficking routes (Paultre et al., 2016), to elucidate genomic and environmental controls over plant phenotypes (Allard et al., 2016; Di Guardo et al., 2017; Montesinos-López et al., 2017; Wang et al., 2019), and to assess gene-to-metabolome associations (Marttinen et al., 2014). In this study, Bayesian model testing validated the supposition that the cellular free VLCFA levels are the most impactful factors affecting product-precursor relationships. Similar relationships between cuticular lipid products and their acyl precursors have been observed in both Arabidopsis and leek. In Arabidopsis, the relationship between VLCFA precursors and alkane products along the developmental gradient from the base to the apex of the leaf was associated with a concomitant increase in *KCS6* expression, a gene encoding for an enzyme required for the synthesis of VLCFA precursors (Busta et al., 2017). Similarly, in leek, VLCFA elongase activity increases along the spatio-temporal gradient of the leaf blade and is correlated with a ∼1000-fold increase in the cuticular ketones, which are the major products in this cuticle (Rhee et al., 1998).

Other factors that significantly impact the cuticular lipid product-precursor relationships include the genetic background and the husk encasement status of the silks. Statistical assessment of the product-precursor ratios suggests that cuticular hydrocarbon production is less active in Mo17 as compared to B73, and that potential environmental cues differentially impact the hydrocarbon-producing pathway between these two genotypes. These different modulators that impact the product-precursor ratios can be mechanistically explored either by transgenic strategies that will evaluate individual genes that could impact product-precursor relationships, or by studying different genotypes that could evaluate the impact of different combinations of alleles on product-precursor relationships.

A key observation made in this study is that the product-precursor ratio increases with increasing alkyl chain length, demonstrating a preference toward the production of longer-chain hydrocarbons via parallel reduction-decarbonylation reactions. There are a number of potential mechanisms that could generate this change between the product-precursor ratio and acyl chain length. These include variations in the substrate preferences of 1) the VLCFA elongation system; 2) the reductase that converts VLCFAs to aldehydes; and 3) the decarbonylase that converts aldehydes to hydrocarbons. Indeed, chain length preferences have been identified for several of the 21 ketoacyl-CoA synthetases (KCSs) responsible for the condensation reaction of the fatty acid elongase that generates VLCFAs of Arabidopsis (Millar and Kunst, 1997; Fiebig et al., 2000; Blacklock and Jaworski, 2006; Joubès et al., 2008; Kim et al., 2013; Hegebarth et al., 2016; Hegebarth et al., 2017) and similar preferences are likely to exist amongst the 26 putative KCSs identified in maize (Campbell et al., 2019). Moreover, the observed substrate specificities of the alkane-forming reductase-decarbonylase enzyme complexes may be associated with homologs of the Arabidopsis CER1 and CER1-LIKE1 proteins, which can each form a unique reductase-decarbonylase complex with CER3 homologs (Pascal et al., 2019). These hypotheses could be tested via heterologous expression of candidate genes within the recently engineered yeast system expressing the maize fatty acid elongase complex (Campbell et al., 2019; Pascal et al., 2019).

## CONCLUSIONS

In this study we demonstrate that the combination of spatio-temporal profiling and multivariate analyses is an effective approach for assessing and comparing metabolic networks that are differentially impacted by developmental and environmental perturbations among different genotypes. Compositional dynamics of this cuticular lipid metabolome were most impacted by silk microenvironment and genotype, with very little impact of silk development. This study establishes that product-precursor ratio dynamics for the hydrocarbon-producing pathway can be inferred from cuticular lipid composition. These product-precursor relationships are complex and dependent upon the composition of cellular VLCFA pools, the genetic background of the organ or tissue, and most significantly, precursor chain length, such that longer chain VLCFAs are preferentially utilized as precursors in hydrocarbon biosynthesis. These findings provide the foundation for the dissection of the underlying mechanisms of the hydrocarbon-producing pathway in response to changes in microenvironment, ultimately leading to a genotype-metabolite-phenotype understanding (Yandeau-Nelson, 2015) of cuticular lipid accumulation, and resultant protective capacity of cuticular lipids on plant surfaces.

## MATERIALS AND METHODS

### Nomenclature

Cuticular lipid metabolites are represented using a *n:y(z)* notation, where *n* represents the number of carbon atoms, *y* the number of double bonds, and *z* the positions of the double bonds in the alkyl chain (Howard and Lord, 2003). This study reports three classes of cuticular lipids; very-long-chain fatty acids (VLCFAs) that are saturated (*FA_n:0_*) or unsaturated (*FA_n:y(z)_*), saturated aldehydes (*Ald_n:0_*), and hydrocarbons (HCs), which include saturated alkanes (*HC_n:0_*), and two classes of alkenes: monoenes (*HC_n:1(z)_*), and dienes (*HC_n:2(z1, z2)_*).

### Plant growth, sample collection and processing

Maize inbred lines B73 and Mo17 and their reciprocal hybrids, B73×Mo17 and Mo17×B73 were grown to maturity at the Iowa State University Agronomy Research Farm (Boone, IA) during the 2014 and 2015 growing years using standard cultivation practices and no supplemental irrigation. Developing ear shoots were covered prior to silking to prevent pollination. Ears were harvested three days after silks had emerged from encasing husk leaves, at a point at which half of the plants in a row were silking (i.e. the mid-silk cohort). Ears were harvested between 11am and 1pm each day and were transported to the laboratory at ambient temperature using insulated chests. The number of biological replicates sampled per genotype were 8 and 12 in the 2014 and 2015 growing years, respectively. Each biological replicate consisted of silks from two ears of similar size and appearance, harvested on the same day. Silk samples were prepared as described in McNinch *et al*. (2020). For each biosample, one sub-sample was flash-frozen in liquid nitrogen and reserved for subsequent total VLCFA profiling and the second sub-sample was immediately subjected to cuticular lipid extraction.

### Lipid extraction and derivatization

Extracellular cuticular lipids were extracted from fresh silks by immersion for 4 min in 9:1 hexanes: diethyl ether supplemented with the internal standards eicosane (1µg/ml), nonadecanoic acid (1µg/ml) and heptadecanol (1µg/ml), as previously described (Loneman, 2017). Extracts were concentrated under a stream of N_2_ gas in a N-EVAP nitrogen evaporator (Organomation Associates, Inc., MA). Extracts were chemically derivatized via transmethylation followed by silylation, as previously described (Loneman, 2017). For total lipid analyses, intra- and extracellular lipids were extracted as previously described (Okazaki, 2015) from 5-8 mg of lyophilized silk tissue that had been pulverized to a fine powder using a Mixer Mill 301 (Retsch GmbH, Germany).

### Chromatography

Gas chromatography of cuticular lipid extracts was performed with an HP-5MS cross-linked (5%) diphenyl (95%) dimethyl polysiloxane column (30m in length; 0.25-mm inner diameter) using helium as the carrier gas, and an Agilent Technologies series 6890 gas chromatograph, equipped with a model 5973 mass detector (Agilent Technologies, Santa Clara, CA). Extracts were introduced to the gas chromatograph via splitless injection of a 2-µl sample and the oven temperature program was as listed for samples that were both transmethylated and silylated (Loneman, 2017). Quantification analysis was performed using the AMDIS software package (Stein, 1999) with assistance from the NIST Mass Spectral library (http://webbook.nist.gov/chemistry/) for compound identification.

Total lipid analysis was performed on a random selection of four biological replicates from the total number of replicates collected in the 2015 growing year. Total lipid extracts were analyzed via liquid chromatography quadrupole time-of-flight mass spectrometry on a Waters Xevo G2 Q-TOF MS combined with a Waters ACQUITY UPLC system in both positive and negative ion modes, as previously described (Kimbara et al.,2013). LC-MS data were recorded using MassLynx4.1 software (Waters) and processed using MarkerLynx XS software (Waters). The data matrices were queried against an in-house lipid library (RIKEN, Japan). For data normalization, the original peak intensity values were divided by the internal standard peak intensity value at m/z 566.382 [M + H]^+^ and m/z 550.351 [M-CH_3_]^−^ for the positive and negative ion modes, respectively.

### Quantitative methods

Cuticular lipid metabolite concentrations were initially calculated relative to the compatible internal standard (i.e. hydrocarbons quantified relative to eicosane, VLCFAs relative to nonadecanoic acid, and alcohols and aldehydes relative to heptadecanol). Because quantification of each metabolite class did not differ depending on choice of standard, all subsequent quantifications were relative to the eicosane standard. Quantification was performed relative to silk dry weight, to avoid potential differences in metabolite concentrations due solely to differences in water content between genotypes or among silk sections. Representative measurements of silk dry weight were calculated for each silk section (A-E) from surrogate ears harvested on the same days as those used for cuticular lipid profiling, and a ratio of dry weight to fresh weight was calculated as a conversion factor to estimate the dry weight of extracted silk samples, as described by Loneman et al. (2017). The measured water content of silk samples was approximately 90% regardless of the position along the silk length, the genetic background, or the growing year.

Detection limits were determined as previously described (Dennison et al., 2019). The detection limits for the datasets collected in growing years 2014 and 2015 were 0.005 and 0.006 µmol g^-1^ dry weight, respectively. Metabolite abundances were censored (Newman et al., 1989) such that concentrations below the detection limit (DL) were assigned a value of DL/2, and concentrations for metabolites that were not detected in a specific condition were assigned the value, DL/10. Consistently low abundance metabolites that were assigned DL/2 or DL/10 values for more than 90% of the samples were removed from the dataset.

### Statistical Methods

The raw metabolite data were log-transformed and pareto-scaled. ANOVA was performed to evaluate the effects of genotype, position along the silk, growing year, and cellular free VCLFA and cellular lipid-associated VLCFA content on cuticular lipid composition. For each genotype, the accumulation dynamics along the silk length for each lipid class were modeled using linear and polynomial (i.e., quadratic, cubic and quartic models) regression. The regression model that best fitted the concentration data for each lipid class was determined by comparing the adjusted R^2^ of the four regression models. Tukey’s HSD tests were applied for post-hoc pairwise comparisons between silk sections either within or among genotypes. These analyses were conducted using the R/stats base package (R core team 2016).

PCA was performed using the prcomp() function in the R/stats package and 95% confidence ellipses were constructed using R/car dataEllipse() function (Fox and Weisberg, 2011). Partial least squares-discriminant analysis (PLS-DA) was performed using R/ropls opls() function that also determined the optimal number of components for the PLS-DA model using seven-fold cross validation (Thevenot et al., 2015). Variable Importance in Projection (VIP) scores were calculated by R/ropls getVipVn() function as a cumulative measure of the contribution of each metabolite in distinguishing among genotypes or silk sections (Pérez-Enciso and Tenenhaus 2003). Individual metabolites with VIP scores >1.0 (i.e. with above-average contribution in sample classification) were deemed as metabolite biomarkers that discriminate between classes. The explained fraction of data variance by a PLS-DA model for explanatory and response variables were reported as R^2^X and R^2^Y, respectively, and the predictive accuracy of the model was reported as Q^2^Y. The response variable for PLS-DA is categorical and therefore transformed into a binarized numerical variable for model construction, and R^2^Y represents the explained variation within the transformed data.

Non-parametric Spearman correlations between every pair of metabolites across silk sections A to E were calculated in each genotype, and were compared among genotypes according to Choi and Kendziorski (2009), with modifications that are described in Supplemental Methods. The metabolite correlation networks were constructed based on topological overlap matrices (TOMs) derived from the correlation matrices using the R/WGCNA package (Langfelder and Horvath, 2008). Briefly, for each genotype, metabolite clusters were initially determined according to TOMs by hierarchical clustering and then pruned by four independent R functions: pam(), cutreeStatic(), and two cutreeDynamic()s with the “methods” argument specified as either “tree” or “hybrid” (Horvath and Langfelder, 2011).

The ratios of *HC_2n-1:0_* :*FA_2n:0_* relative to the carbon chain length *2n* were fitted into a linear regression model as the response variable using the R/stat function lm(), with precursor chain length, genotype, silk encasement status, growing year, and the corresponding interaction terms as the explanatory variables. The prediction interval for regression was calculated by R/stat function predict(). Bayesian Model Comparison, using the R/BayesFactor package (Morey and Rouder 2015), was applied to assess the association of the product-precursor ratio with these explanatory variables and two additional explanatory variables (concentration of cellular free VLCFAs and concentration of lipid-associated VLCFAs). Bayesian model construction and comparison are described in detail in Supplemental Methods.

**Supplemental methods**: Supplemental Statistical Methods

**Supplemental Figure S1**: Accumulation of cuticular monoenes along the spatio-temporal gradient of silks.

**Supplemental** Figure S2: Correlation-based clustering of silk cuticular lipid abundance data for hybrids, B73×Mo17 and Mo17×B73.

**Supplemental Figure S3**: Clustering of silk samples by partial least squares discriminant analysis (PLS-DA) based on different genotypes and different silk sections.

**Supplemental Figure S4**: The spatio-temporal accumulation patterns of hydrocarbon (*HC_2n-1:0_*) and VLCFA (*FA_2n:0_*) product-precursor pairs in inbreds B73 and Mo17.

**Supplemental Figure S5**: Accumulation of cuticular hydrocarbon products and corresponding cuticular VLCFAs, and the regression of the product-precursor ratios on chain length for hybrids, B73×Mo17 and Mo17×B73.

**Supplemental Table S1**: Spatio-temporal profiles of cuticular lipids from maize silks of B73, Mo17, and the reciprocal hybrids that were collected in the 2014 and 2015 growing years.

**Supplemental Table S2**: Comparison of four polynomial regression models that depict the dynamics of cuticular metabolite accumulation levels among the silk sections in the indicated maize lines.

**Supplemental Table S3**: Two-way and two-way nested ANOVAs that test the main effect of genotype (G), silk section (S) and G × S interaction on each cuticular lipid metabolite.

**Supplemental Table S4**: Two-way and three-way ANOVAs of total cuticular lipids and individual lipid metabolite classes.

**Supplemental Table S5**: Variable importance in projection (VIP) score for each lipid in two PLS-DA models built to discriminate genotypes or silk sections based on the cuticular lipid metabolome compositions.

**Supplemental Table S6**: Spatio-temporal profiles of cellular VLCFAs from maize silks of B73 and Mo17 at 2015 growing year.

**Supplemental Table S7**: Pearson correlations between cellular fatty acids (free VLCFAs and lipid-associated VLCFAs) and cuticular VLCFAs at individual acyl chain lengths.

**Supplemental Table S8**: Identification of the optimal VLCFA Bayesian regression model that incorporated cellular free VLCFAs and/or cellular lipid-associated VLCFAs as predictors.

## ACKNOWLEDGEMENTS

The authors thank Drs. Ann Perera, Lucas Showman, and Zhihong Song at the Iowa State University’s W. M. Keck Metabolomics Research Laboratory for assistance and advice regarding GC-MS analysis. We thank Pauline Aamodt, Grace Kuehne, Sarah (Weirich) Hennings, and Kyle King for assistance with sample collection and processing.

